# Accounting for point count ambient noise increases population size estimates

**DOI:** 10.1101/2021.04.06.438644

**Authors:** Emily K. Anderson, Pete F. Kerby-Miller, Julia M. Pupko, Nathaniel R. Sharp, Kevin S. Tolan, Jason M. Hill

## Abstract

Ambient noise is an integral component of natural environments, but it also creates challenges for avian monitoring programs. Ambient noise can mask bird vocalizations from observers during point counts, which may lead to systematic undercounting of birds in noisy environments. Here, we estimate detection probability and population size in models that either account for or omit the influence of ambient noise. We used data for four bird species, from 2228 point counts that were conducted during the 2019 Mountain Birdwatch field season. Community scientists assessed ambient noise using a simple scale. Despite relatively quiet conditions at sampling locations (<u>*x*</u> = 2.48), our information theoretical approach favored N-mixture models that incorporated ambient noise into the detection function for all four species. At the noisiest sampling locations, our models predicted detection probabilities that were as low as 10% for some species. Accounting for ambient noise resulted in a modest, mean increase of ≤3.29% in the total population size for each species. Following our approach, other researchers can easily incorporate ambient noise assessments into their field protocols and analyses, with minimal costs or added complexity, to increase the comparability of studies conducted within different acoustic environments.

---

Ambient noise is a constant presence in natural environments, and an integral part of an ecosystem’s soundscape: the complex arrangement of sounds from biological, geophysical, and anthropogenic sources that create acoustical patterns in space and time (Pijanowski et al. 2011). Ambient noise is the portion of the soundscape that is uninformative to the listener. Birds that vocally communicate must compete with local ambient noise to be heard (Pijanowski et al. 2011). In the presence of elevated ambient noise, birds may modify the volume (Wood and Yezerinac 2009), frequency (Bermúdez-Cuamatzin et al. 2011; Gallardo Cruz et al. 2020), or timing of their vocalizations and other audible signaling to mitigate masking (Gil et al. 2015).

Ambient noise also imposes substantial challenges for bird monitoring regimes. Ambient noise can negatively affect the ability of observers to detect, localize, and identify birds during point counts (Simons et al. 2007, 2009; Koper et al. 2016). Furthermore, the influence of ambient noise on detectability and availability may vary between species (e.g., based on differences in vocal behavior) and with the specific identity of the ambient noise source (Koper et al. 2016; Yip et al. 2017). These research findings have been largely drawn from studies using simulation and playback experiments under controlled conditions, whereas avian field studies often limit surveying to conditions without excessive ambient noise (Huff et al. 2000). Nonetheless, the implications of not accounting for ambient noise in bird monitoring regimes are potentially consequential. Failure to account for ambient noise in the analysis of bird count data could lead to distorted inference about local density, habitat preferences, population trends, and species richness and community structure (*sensu* Schmidt 2005; Kéry and Royle 2009).

These concerns are relevant to the analysis and interpretation of data from Mountain Birdwatch, a long-term community science project that annually monitors 10 high-elevation breeding bird species along remote hiking trails in the northeastern United States. Due to the steep, rugged terrain, hiking trails in our study region are often located along the relatively gentle gradient within ravines or switch-backed along ridgelines. These locations are often subject to undesirable amounts of ambient noise, particularly from wind and ephemeral streams; occasional ambient noises stemming from logging equipment, all-terrain vehicles, aircraft, and nearby hiking groups with barking dogs are also present (JMH pers. obs.). Observers use meteorological forecasts to target their surveys during periods of fair weather. Unlike weather, however, the ambient noise at sampling locations is largely unpredictable, and is a source of frustration for a varying number of Mountain Birdwatch community scientists each year (JMH pers. obs.).

Here, we used N-mixture models to investigate the relationship between ambient noise, detection probability, and resulting local population size estimates for four montane bird species using >2000 point counts from >500 sampling locations. We purposefully chose species with a wide range of vocal complexity, singing frequency, and song volume. We used a model selection approach to determine if ambient noise (measured on a simple 1-10 scale) helped account for unexplained variation in detection probability. We hypothesized that ambient noise levels would negatively relate to detection probability. To quantify the potential consequences of omitting ambient noise from our analyses, we compared species-specific population estimates generated from models that included or omitted the ambient noise detection covariate. We hypothesized that inclusion of the ambient noise covariate would account for individuals otherwise masked by relatively loud ambient sounds, and thus result in larger population size estimates. Based on our results, we make study design recommendations and specific modeling recommendations for incorporating ambient noise into future analyses.

## METHODS

Mountain Birdwatch is a community science effort to monitor the breeding populations of 10 montane bird species throughout the northeastern United States (Hill and Lloyd 2017). Initiated in 2000, community scientists annually conduct avian sampling at up to 747 long-term sampling locations located along 129 routes. Sampling locations (≤6 per route) are ≥0.25 km apart and primarily located along hiking trails in the spruce-fir forest. Individual sampling locations range in elevation from 582 to 1502 m and stretch from the Catskill Mountains, NY (41.9° N latitude) to northern Maine (46.0° N latitude).

At each sampling location, a lone observer conducts four independent five-minute unlimited distance point counts in succession. All sampling locations on a route are annually surveyed during a single day in June (e.g., 96% in 2019) or (rarely) early July. The first point count survey on a route begins 0.75 hr before sunrise. Counts are conducted during fair weather (wind speeds <19 kph, >0°C, and no rain), and the vast majority (98.92%) of avian detections are made aurally (JMH unpub. data).

In 2019, we encouraged Mountain Birdwatch observers to quantify the intensity of ambient noise levels at each sampling location using an intensity scale from 1 (“virtually no ambient noise”) to 10 (“very loud ambient noise levels that likely prevent hearing avian vocalizations if present”). We asked observers to estimate the mean ambient noise level across the entire 20-minutes of counts at a sampling location, because the most common ambient noise sources (wind and flowing water) at these locations tend to be fairly constant and not episodic (JMH pers. obs.). Previously, we experimented with assessing ambient noise via a less subjective method, a smartphone app, but we abandoned that idea after field tests verified that the app confounded ambient noise and avian vocalizations.

### Bird species selection rationale

We *a priori* chose four Mountain Birdwatch species with a diversity of song complexity and vocal behavior. Yellow-bellied Flycatchers (*Empidonax flaviventris*) have a relatively short, simple song that is delivered between 6-10 times per minute (Gross and Lowther 2020) that generally falls between 2 and 6 kHz. Blackpoll Warblers (*Setophaga striata*) songs are a high-pitched trill (8–10 kHz) lasting roughly 2 seconds (Brand 1938). Winter Wren (*Troglodytes hiemalis*) songs are relatively long, 5-10 seconds in length, and highly complex with a wide range of frequencies from below 3 to around 8 kHz (Hejl et al. 2020). White-throated Sparrow (*Zonotrichia albicollis*) songs are relatively simple, consisting of several whistled notes, the frequency of which does not surpass ~6 kHz in New York (Borror and Gunn 1965).

### N-Mixture model selection

We fit N-mixture models in a frequentist framework in R version 4.0.3 (R Core Team 2020) using the *pcount* function from the unmarked package version 1.0.1 (Fiske and Chandler 2011). N-mixture models estimate local abundance N after accounting for imperfect detection *p* by observers (Royle 2004; Kéry and Schaub 2011). We performed model selection using Akaike’s information criterion (AIC; Burnham and Anderson 2002) separately for each species, using the 2019 Mountain Birdwatch data from sampling locations where observers assessed ambient noise.

In addition to the ambient noise covariate, we considered two additional covariates of detection probability (point count start time and survey date) and local abundance (elevation and latitude). These candidate covariates and their quadratic terms were identified as being informative in past modeling efforts using nine years of Mountain Birdwatch data (Hill 2020). For bird species monitored by Mountain Birdwatch, local abundance typically peaks at mid-elevations and mid-latitudes, while detection probability generally declines throughout the morning and the month of June (Hill 2020). Observers recorded the point count start time (decimal hours) and survey date (1 June = 1, 2 June = 2, etc.). We used Zonal Statistics in ArcGIS version 10.6.1 (ESRI 2018) to calculate the elevation (m) and latitude (°N) at each Mountain Birdwatch sampling location. All covariates were grand-mean-centered and scaled by one standard deviation to improve computational efficiency.

We used a systematic 3-step model selection process similar to Hill and Lloyd (2017) and Kidwai et al. (2019). We started with a near-global model (a global model without interactions) that included all detection and local abundance covariates and their quadratic terms, and compared the performance of poisson, zero-inflated poisson, and negative binomial N-mixture models (step 1). While holding the local abundance model components constant, we identified the parsimonious detection model structure by considering all combinations of the detection covariates (step 2). We retained the parsimonious detection model structure and compared all combinations of the local abundance components (step 3). Switching the order of steps 2 and 3 still resulted in the same parsimonious model for each species (JMH unpub. data). Quadratic terms were only included in a model when the lower order polynomial term was included, and we did not consider interactions between covariates. All candidate models and the complete model selection results are available in the Appendix (Supplemental Tables S1:S4). For each species, we assessed model fit of the near-global model with the parboot function in unmarked using 1000 simulations; there was no indication of a lack of fit (all P > 0.05).

### Bayesian formulation of the parsimonious N-Mixture models

Finally, we used WinBUGS version 1.4.3 (Lunn et al. 2000) from within program R to fit the parsimonious model for each species from step 3 in a Bayesian framework to improve model interpretation and assessment (Kéry and Royle 2015). In all models we specified a uniform prior on detection probability (*p* ~ Uniform(0,1)) and weakly informative logistic priors centered at 0 with scale parameter 1 (~ dlogis(0,1)) for intercepts and covariate coefficients modeling local abundance and detection (Northrup and Gerber 2018). For the proportion of suitable locations (ϕ) in zero-inflated poisson N-mixture models, we specified a uniform prior (ϕ ~ Uniform(0,1)) that was subsequently logit-transformed (Kéry and Royle 2015). We conducted a sensitivity analysis on the priors for intercepts and covariate coefficients by comparing the mean and standard errors of parameters produced via 1) packaged unmarked (i.e., maximum likelihood estimates), 2) normally distributed priors with mean zero and a variance of 2, 10, or 20, and 3), our weakly informative logistic priors. All prior choices produced similar parameter and uncertainty estimates.

From the parsimonious model for each species, we retained 3000 total iterations from 3 Markov chain Monte Carlo chains (MCMC), thinned at a rate of 1:100, with a burn-in of 10,000 iterations. For each species’ model results, we examined plots of the residuals and traceplots of the posteriors and verified that Gelman-Rubin statistics 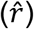 were <1.01 (Gelman and Rubin 1992).

### Contribution of the ambient noise covariate to population size estimates

The ultimate goal of researchers using N-mixture models is often to estimate the total population size (N_total_) of study organisms surrounding the sampling locations (Kéry and Royle 2015). We devised a simple approach to assess the influence of the ambient noise covariate on estimates of N_total_ from the posterior of the Bayesian formulation of the N-mixture models. For those species with the ambient noise covariate in their parsimonious model, we fit that same model but without the ambient noise covariate. From the models’ posteriors (*n* = 3000) for each species, we then compared estimates of total population size generated from the two models with (N_total-with-noise_) and without (N_total-sans-noise_) the ambient noise covariate. We expressed this mean change in total population size as a percent to facilitate comparison between species:

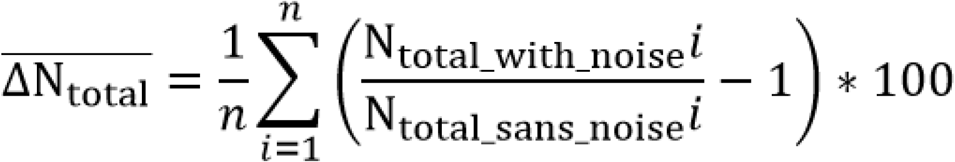

We also undertook a simple approach to assess how the inclusion of ambient noise affected the precision of the mean N_total_ estimate by comparing the coefficient of quartile variation (cqv) estimates of N_total_ from models fitted with and without the ambient noise covariate. The cqv is expressed as a percentage, robust to non-normal distributions, and a useful measure of the relative dispersion of a parameter (Bonett 2006). We generated separate estimates of the coefficient of quartile variation (cqv) for N_total_ from those two models as (*Q*3 − *Q*1)/(*Q*3 + *Q*1), where Q1 and Q3 are the 25th and 75th population percentiles of N_total_, respectively. Comparing cqv estimates of N_total_ from models fitted with and without the ambient noise covariate allowed us to assess how the inclusion of ambient noise affected the precision of N_total_.

Results are presented as mean ± SD (95% credible interval [CRI]) unless otherwise stated. The open‐access data used in this paper are available online (Vermont Center for Ecostudies 2019).

## RESULTS

In 2019, 83 Mountain Birdwatch observers collected ambient noise estimates at 557 sampling locations along 110 routes, and they conducted 2228 5-minute point counts. On average, observers rated ambient noise as 2.48 ± 1.70 [min-max = 1-10] across sampling locations (Fig. 1). Observers scored 6% of sampling locations as having ambient noise ratings ≥5 and detected each of the four species at 33% (Yellow-bellied Flycatcher), 47% (White-throated Sparrow), 55% (Blackpoll Warbler) or 65% (Winter Wren) of sampling locations.

**Figure 1.**
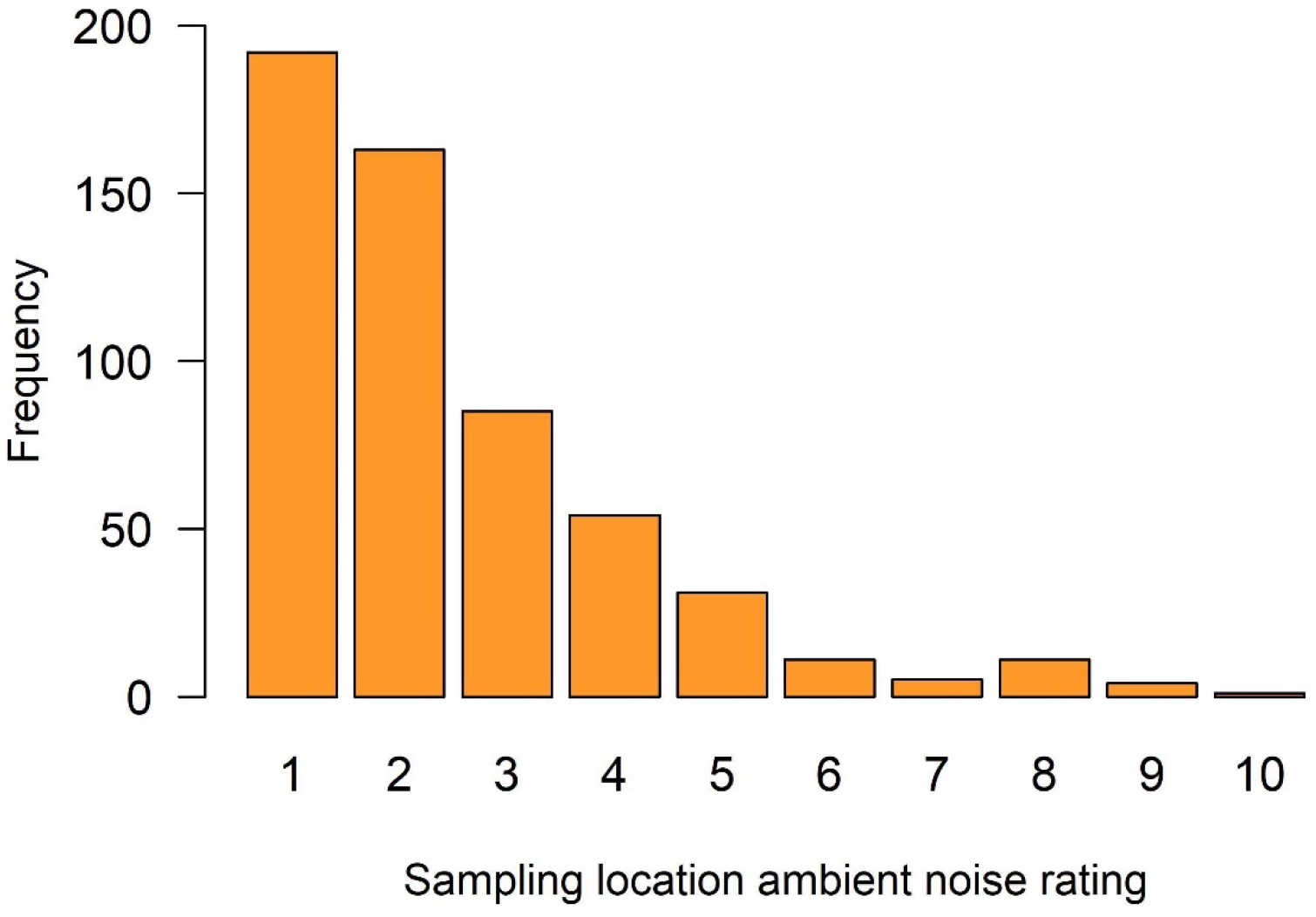
Histogram of ambient noise values (scale of 1 to 10) as assessed by 83 Mountain Birdwatch observers at 557 point count sampling stations in 2019.

Our model selection process favored poisson (for Winter Wren and Blackpoll Warbler) or zero-inflated poisson (for White-throated Sparrow and Yellow-bellied Flycatcher) N-mixture models, and the ambient noise detection covariate was included in the parsimonious model for each species (Table 1; see Appendix for complete model results for each species). For Winter Wren, a one unit increase in the ambient noise scale was associated with a 1.70% (± 0.74%, 0.25% to 3.17%) reduction in detection probability; detection probability varied quadratically with ambient noise levels for the other three species and peaked at 4.27 (Yellow-bellied Flycatcher), 4.00 (Blackpoll Warbler), and 2.64 (White-throated Sparrow; Fig. 2).

**Table 1.** Detection probability (*p*) and local abundance (N) covariates included in the parsimonious model for species, which included linear (X) and quadratic (X^2^) terms.

| Species | $p$ | | N | | |
| --- | --- | --- | --- | --- | --- |
|  | Noise | Count time | Survey date | Elevation | Latitude |
| Winter Wren | X | $X^2$ | | $X^2$ | |
| Blackpoll Warbler | $X^2$ | | X | $X^2$ | X |
| White-throated Sparrow | $X^2$ | $X^2$ | $X^2$ | X | X |
| Yellow-bellied Flycatcher | $X^2$ | | $X^2$ | $X^2$ | $X^2$ |

**Figure 2.**
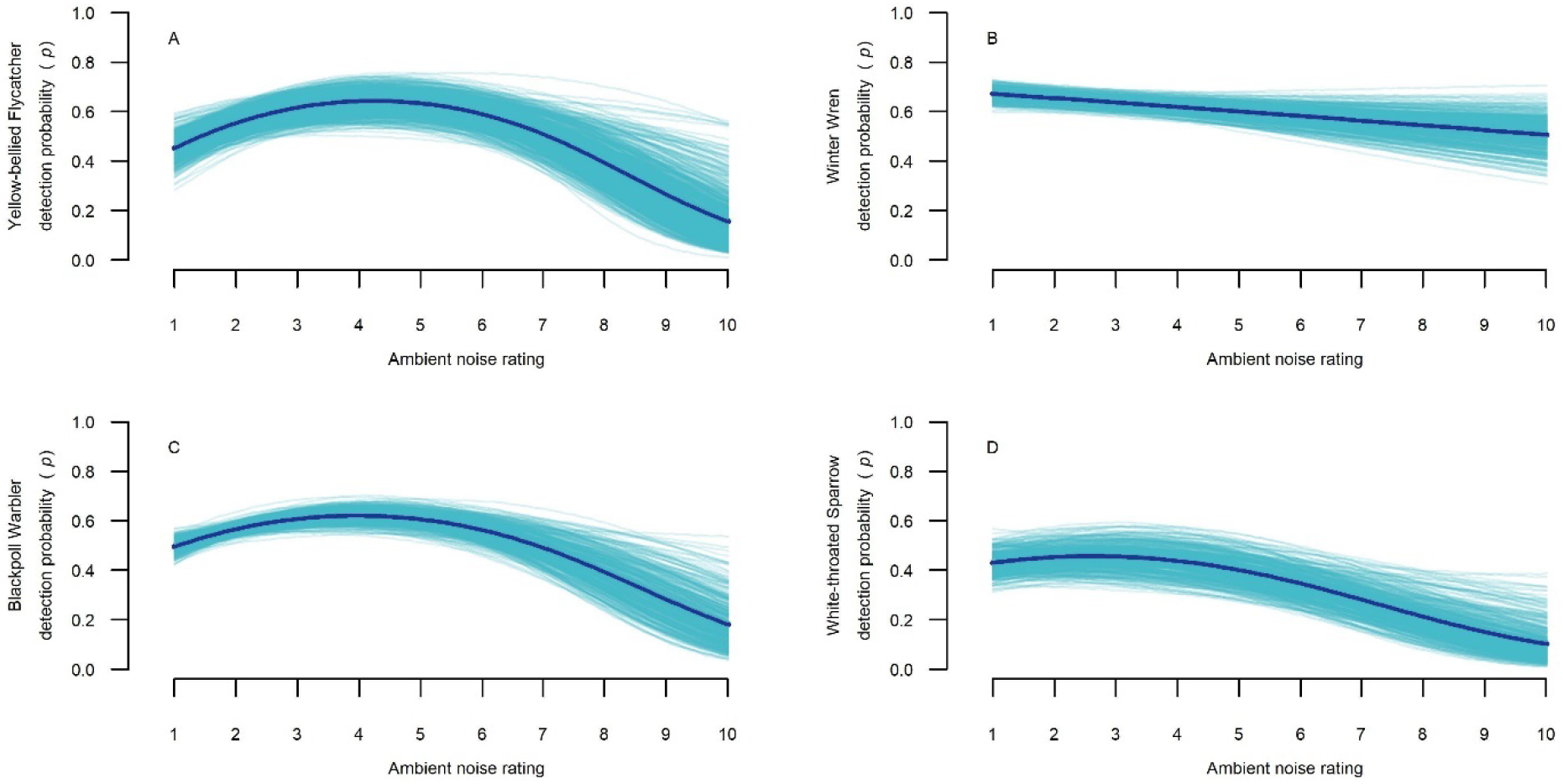
The relationship between background noise and the detection probability of four bird species (panels A-D) monitored at 557 Mountain Birdwatch sampling locations, as assessed via N-mixture models. The darker and thicker line in each panel represents the overall mean predicted model response, while the lighter, thinner lines show 1000 predictions generated from the model posterior.

Compared to the least noisy sampling locations (ambient noise = 1), locations with an ambient noise rating of 10 were associated with a mean reduction in detection probability of 27.58% across species: Winter Wren (16.49% ± 7.45%; 2.25% to 31.57%), Blackpoll Warbler (31.35% ± 8.20%, 10.92% to 43.55%), White-throated Sparrow (32.85% ± 7.58%, 14.03% to 44.33%), and Yellow-bellied Flycatcher (29.61% ± 8.92%, 7.82% to 43.51%).

As we hypothesized, inclusion of the ambient noise covariate resulted in a (modest) increase in N_total_ of ≤3.29% for each species; the 95% CRI for these increases, however, all overlapped 0.00% (Fig. 3). With the inclusion of the ambient noise covariate the cqv of N_total_ became less precise for all species: Winter Wren (increased from 1.12% to 1.20%), Blackpoll Warbler (1.56% to 1.66%), White-throated Sparrow (2.60% to 2.76%), and Yellow-bellied Flycatcher (2.98% to 3.39%).

**Figure 3.**
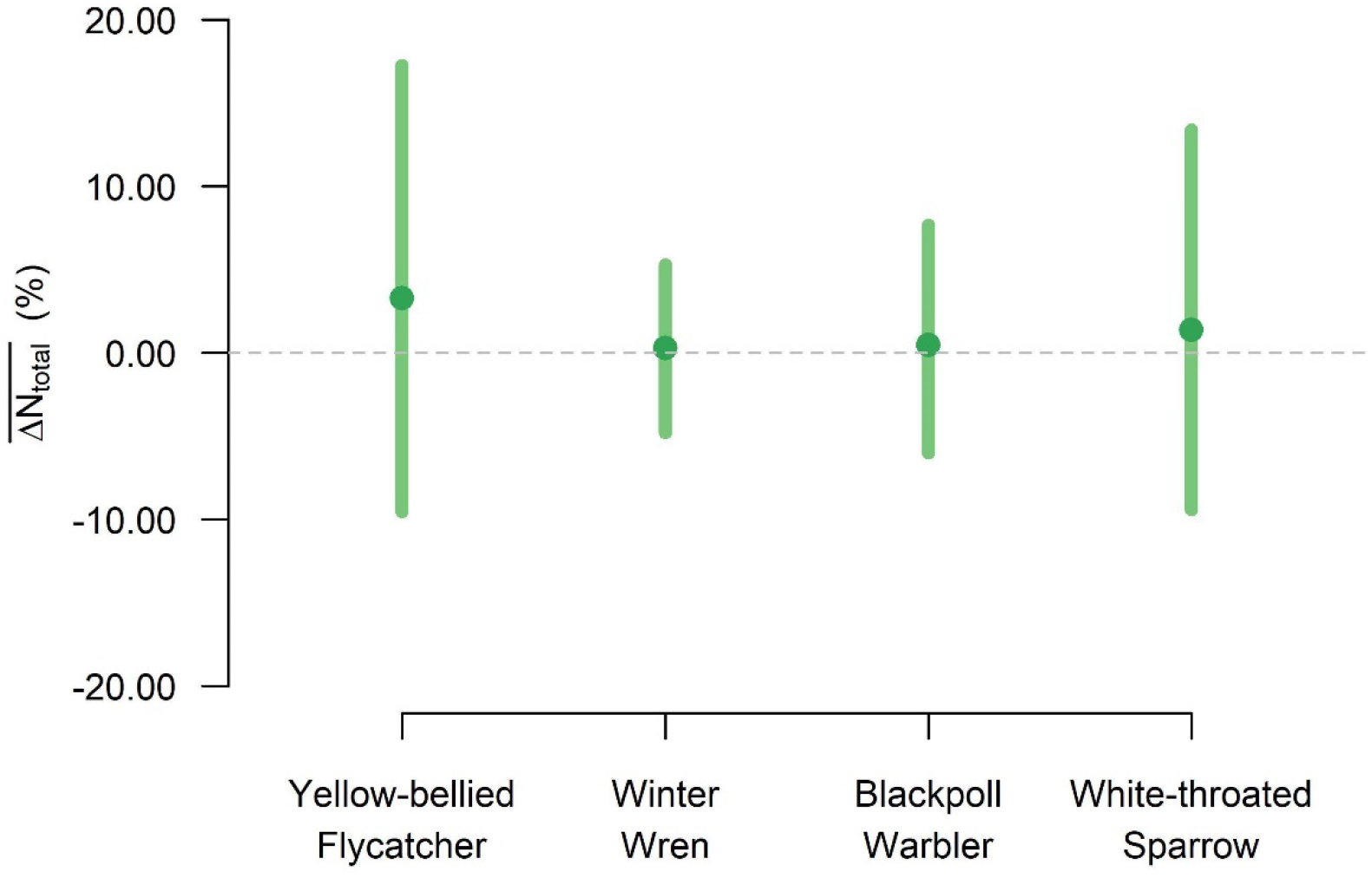
The mean (circles) contribution of the background noise covariate to estimates of total population size 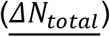 for each of the four Mountain Birdwatch species, expressed as a percent. For example, total population size estimates for Winter Wren were on average 1.39% larger (95% CRI: −9.39%, 13.39%) in models that accounted for ambient noise compared to models that did not account for ambient noise. Inclusion of the background noise covariate resulted in a (modest) increase in N_total_ of ≤3.29% for each species. The 95% CRI (vertical bars) for these increases all overlapped 0.00% dashed gray line).

## DISCUSSION

Ambient noise negatively affected the detection probability of all species included in our analyses, even though the vast majority (94%) of our sampling locations were relatively quiet (ambient noise score ≤4). Our findings support previous experimental simulation and playback results that documented negative ambient noise effects on avian detectability at moderate ambient noise volumes (Simons et al. 2007, 2009; Pacifici et al. 2008). While the bulk of studies that have investigated ambient noise effects on avian detectability used simulated bird vocalizations or artificial ambient noise (i.e., Alldredge et al. 2007; Yip et al. 2017), ours is distinct in examining both naturally-occurring bird vocalizations and naturally-generated ambient noise.

The detection probabilities of all species in our study were inversely related to ambient noise, generally (Fig. 2). These relationships are illustrated in an R Shiny app that we developed to approximate field conditions (https://pkmkp.shinyapps.io/mbw_audio/). Overall, the relationships between detection probability and ambient noise were quite similar for three species (Panels A, C, and D; Fig. 2). Winter Wren, however, had the smallest overall decrease in mean detection probability and was the only species whose detection probability monotonically declined with increasing ambient noise (Panel B; Fig. 2). The Winter Wren song characteristics that might have contributed to these results include its wide range of frequencies, relatively long length, and comparatively loud volume (McCallum 2005; Hejl et al. 2020). Winter Wren songs can encompass over 300 notes and 40 different syllables per song (Kroodsma 1980) and are “ten times louder than a crowing rooster” (Brackenbury 1982). Some experimental evidence indicates that species whose songs encompass many frequencies (i.e., are more complex) may have a greater chance of detection during point counts than species whose calls vary less, especially in the presence of noise interference (Koper et al. 2016).

Amplitude (volume) of ambient noise has a stronger masking effect on bird songs than ambient noise frequency (Apol et al. 2019). Thus, the combination of the high complexity and relative loudness of Winter Wren songs makes it unlikely that all song frequencies would be effectively masked by ambient noise. Furthermore, the Winter Wren’s relatively long song length could provide greater opportunity for detection by observers (McCallum 2005). Although the shape of the relationship between ambient noise and Winter Wren detection probability differed from the other three focal species (Fig. 2), our cumulative results highlight the importance of considering ambient noise effects within avian monitoring schemes for any species.

Across all four focal species, detection probability was not 0 at an ambient noise level of 10, contrary to expectations (Fig. 2). Most of our sampling sites were relatively quiet and observers rarely reported high levels of ambient noise, possibly contributing to the non-zero detection probability for all species. We had previously adjusted sampling locations due to volunteer feedback: prior to 2019, several participants reported persistent high levels of ambient noise that considerably obscured avian vocalizations at some locations, leading us to permanently retire some two dozen sampling locations. Despite our relatively quiet sampling locations, incorporating the ambient noise covariate still improved the performance of our models across all species. The improvements seen in our models’ performance suggests that including ambient noise is important when modelling population estimates, even if ambient noise levels are low. Our findings also suggest that researchers modeling bird populations in locations subject to higher levels of background noise could improve the accuracy of their total population estimates by using a similar approach.

Our results are consequential for the community scientists who collect data for Mountain Birdwatch each year. Frustratingly high levels of ambient noise can lead to increased observer attrition rates, and a subsequent under sampling of relatively noisy sampling locations (Marsh and Cosentino 2019). In previous years, Mountain Birdwatch observers whose sites were especially noisy expressed concern that their data would not provide a valuable contribution to our study, due to the level of noise interference (JMH pers. obs.). In our study, population estimates from models that included ambient noise were higher than those without, indicating that some birds were not detected by observers due to ambient noise (Fig. 3). These results show that the variation in population estimates attributed to ambient noise interference can be accounted for with simple additions to study design, potentially reassuring future community scientists that their data is valuable, despite noisy conditions. Furthermore, our approach to noise quantification is easily transferable.

Numerous opportunities exist for future studies to build upon our research. We selected a diverse assemblage of species within the Mountain Birdwatch dataset, but survey locations were predominantly located within a montane conifer biome. Future studies should test whether the relationship between detection probability and ambient noise remains consistent across varying acoustic environments. Our study utilized a simple scale for rating ambient noise isolated from bird vocalizations, but future studies may want to explore less-subjective instantaneous digital measurements of ambient noise when no birds are vocalizing. Another opportunity for future study is the extent to which different ambient noise sources – such as wind, water, and motorized vehicles – affect detection probability. We concluded that N_total_ increased when we accounted for ambient noise, as did the uncertainty associated with the N_total_ estimate. However, this may not be consistent under all conditions and it is worth exploring whether changing other factors, such as increasing the proportion of noisy study sites, may reduce uncertainty. To facilitate these efforts, we have suggested informed priors for the covariates included in our study for use in future Bayesian analyses (Supplemental Table S9).

## Supporting information

Supplemental Table 0 Definitions

## ACKNOWLEDGEMENTS

Mountain Birdwatch is conducted entirely within unceded territories of the Munsee Lenape and nations of the Haudenosaunee and Wabanaki Confederacies. Survey sites are located within the homeland of the Abenaki, Pequawket, Arosaguntacook, Nanrantsouak, and Penobscot nations of the Wabanaki Confederacy and within that of the Kanien’kehá:ka nation of the Haudenosaunee Confederacy. We would like to thank those who have facilitated access for Mountain Birdwatch surveys, and all of the community scientists (https://mountainbirds.vtecostudies.org/the-community-scientists/) whose tireless efforts have made this work possible. Funding for portions of this work were generously provided by the Charles E. and Edna T. Brundage Foundation and the Robert F. Schumann Foundation.

