## Supplemental Table 0 Definitions for "Accounting for point count ambient noise increases population size estimates"

| Tables | Description |
|---|---|
| S1-S4 | Complete N-mixture model selection results for Yellow-bellied Flycatcher (Table S1), Winter Wren (Table S2), Blackpoll Warbler (Table S3), and White-throated Sparrow (Table S4). |
| S5-S8 | Parameter estimates from the parsimonious model for each species: Yellow-bellied Flycatcher (Table S5), Winter Wren (Table S6), Blackpoll Warbler (Table S7), and White-throated Sparrow (Table S8). |
| S9 | Composite parameter estimates to be considered for informed priors in subsequent Bayesian analyses. We generated these estimates from the posteriors of the final models. We created composite parameter estimates separately from those models with either univariate or quadratic forms of each covariate. |

| Abbreviation or symbol used in the supplementary tables. | Definition |
|---|---|
| N | Local abundance |
| p | Detection probability |
| Φ | Proportion of suitable sites (only applicable in Poisson zero-inflated models) |
| PZIP | Zero-inflated poisson distribution |
| PN | Poisson distribution |
| NB | Negitive binomial distribution |
| K | Number of model parameters. All models have global intercept for detection probability and local abundance. Zero-inflated poisson and negative binomial models contain one additional base parameter than a poisson distribution. |
| AIC | Akaike's Information Criterion |
| ΔAIC | Delta AIC: the change in AIC score between the model being compared and the parsimonious model |
| AIC.Wt | AIC weight: proportion of the amount of predictive power provided by the full set of models being compared |
| Cum.Wt | Cummulative AIC weight |
| LL | Log-likelihood |
| Date | Survey date (1 June =1, 2 June =2, etc.) |
| Date2 | Survey date, quadratic term |
| Count.Time | Survey start time (decimal hours) |
| Count.Time2 | Survey start time, quadratic term |
| Noise | Background noise (recorded on a scale of 1-10) |
| Noise2 | Background noise, quadratic term |
| Elevation | Elevation (m) |
| Elevation2 | Elevation, quadratic term |
| Latitude | Latitude (˚N) |
| Latitude2 | Latitude, quadratic term |
| β | Mean parameter estimate |
| SD | Standared deviation of the parameter estimate |
| LCRI | Lower bound, 95% credible interval |
| UCRI | Upper bound, 95% credible interval |
| totalN | Total population size (derived parameter) |
| mean.detection | Overall mean detection across all sampling locations (derived parameter) |
