## Supplemental Table 1 for "Accounting for point count ambient noise increases population size estimates"

Supplementary Table S1

Yellow-bellied Flycatcher--model selection step 1:

| Candidate model | Distribution | K | AIC | ΔAIC | AIC.Wt | Cum.Wt | LL |
|---|---|---|---|---|---|---|---|
| p(Count.Time + Count.Time2 + JDAY + JDAY2 + Noise + Noise2), N(Elevation + Elevation2 + Latitude + Latitude2) | PZIP | 13 | 2037.04 | 0.00 | 0.53 | 0.53 | -1005.52 |
| p(Count.Time + Count.Time2 + JDAY + JDAY2 + Noise + Noise2), N(Elevation + Elevation2 + Latitude + Latitude2) | NB | 13 | 2037.31 | 0.27 | 0.46 | 0.99 | -1005.65 |
| p(Count.Time + Count.Time2 + JDAY + JDAY2 + Noise + Noise2), N(Elevation + Elevation2 + Latitude + Latitude2) | PN | 12 | 2045.04 | 8.00 | 0.01 | 1.00 | -1010.52 |

Model selection step 2:

| Candidate model | Distribution | K | AIC | ΔAIC | AIC.Wt | Cum.Wt | LL |
|---|---|---|---|---|---|---|---|
| p(Count.Time + Date + Date2 + Noise + Noise2), N(Elevation + Elevation2 + Latitude + Latitude2) | PZIP | 12 | 2035.16 | 0.00 | 0.58 | 0.58 | -1005.58 |
| p(Count.Time + Count.Time2 + JDAY + JDAY2 + Noise + Noise2), N(Elevation + Elevation2 + Latitude + Latitude2) | PZIP | 13 | 2037.04 | 1.88 | 0.22 | 0.80 | -1005.52 |
| p(Date + Date2 + Noise + Noise2), N(Elevation + Elevation2 + Latitude + Latitude2) | PZIP | 11 | 2037.48 | 2.32 | 0.18 | 0.98 | -1007.74 |
| p(Count.Time + Date + Date2), N(Elevation + Elevation2 + Latitude + Latitude2) | PZIP | 10 | 2044.95 | 9.79 | 0.00 | 0.98 | -1012.47 |
| p(Date + Date2), N(Elevation + Elevation2 + Latitude + Latitude2) | PZIP | 9 | 2045.82 | 10.67 | 0.00 | 0.99 | -1013.91 |
| p(Count.Time + Date + Date2 + Noise), N(Elevation + Elevation2 + Latitude + Latitude2) | PZIP | 11 | 2046.16 | 11.00 | 0.00 | 0.99 | -1012.08 |
| p(Count.Time + Noise + Noise2), N(Elevation + Elevation2 + Latitude + Latitude2) | PZIP | 10 | 2046.39 | 11.24 | 0.00 | 0.99 | -1013.20 |
| p(Date + Date2 + Noise), N(Elevation + Elevation2 + Latitude + Latitude2) | PZIP | 10 | 2046.83 | 11.67 | 0.00 | 0.99 | -1013.42 |
| p(Count.Time + Date + Noise + Noise2), N(Elevation + Elevation2 + Latitude + Latitude2) | PZIP | 11 | 2046.86 | 11.70 | 0.00 | 0.99 | -1012.43 |
| p(Count.Time + Count.Time2 + Date + Date2), N(Elevation + Elevation2 + Latitude + Latitude2) | PZIP | 11 | 2046.92 | 11.76 | 0.00 | 1.00 | -1012.46 |
| p(Count.Time + Count.Time2 + Noise + Noise2), N(Elevation + Elevation2 + Latitude + Latitude2) | PZIP | 11 | 2047.94 | 12.79 | 0.00 | 1.00 | -1012.97 |
| p(Count.Time + Count.Time2 + Date + Date2 + Noise), N(Elevation + Elevation2 + Latitude + Latitude2) | PZIP | 12 | 2048.14 | 12.98 | 0.00 | 1.00 | -1012.07 |
| p(Count.Time + Count.Time2 + Date + Noise + Noise2), N(Elevation + Elevation2 + Latitude + Latitude2) | PZIP | 12 | 2048.50 | 13.34 | 0.00 | 1.00 | -1012.25 |
| p(Noise + Noise2), N(Elevation + Elevation2 + Latitude + Latitude2) | PZIP | 9 | 2049.30 | 14.14 | 0.00 | 1.00 | -1015.65 |
| p(Date + Noise + Noise2), N(Elevation + Elevation2 + Latitude + Latitude2) | PZIP | 10 | 2050.41 | 15.25 | 0.00 | 1.00 | -1015.21 |
| p(Count.Time), N(Elevation + Elevation2 + Latitude + Latitude2) | PZIP | 8 | 2051.75 | 16.59 | 0.00 | 1.00 | -1017.87 |
| p(Count.Time + Date), N(Elevation + Elevation2 + Latitude + Latitude2) | PZIP | 9 | 2052.77 | 17.62 | 0.00 | 1.00 | -1017.39 |
| p(Count.Time + Noise), N(Elevation + Elevation2 + Latitude + Latitude2) | PZIP | 9 | 2052.86 | 17.70 | 0.00 | 1.00 | -1017.43 |
| p(Count.Time + Count.Time2), N(Elevation + Elevation2 + Latitude + Latitude2) | PZIP | 9 | 2053.70 | 18.55 | 0.00 | 1.00 | -1017.85 |
| p(Count.Time + Date + Noise), N(Elevation + Elevation2 + Latitude + Latitude2) | PZIP | 10 | 2053.79 | 18.63 | 0.00 | 1.00 | -1016.89 |
| p(Noise), N(Elevation + Elevation2 + Latitude + Latitude2) | PZIP | 8 | 2054.07 | 18.91 | 0.00 | 1.00 | -1019.03 |
| p(Date), N(Elevation + Elevation2 + Latitude + Latitude2) | PZIP | 8 | 2054.69 | 19.53 | 0.00 | 1.00 | -1019.34 |
| p(Count.Time + Count.Time2 + Date), N(Elevation + Elevation2 + Latitude + Latitude2) | PZIP | 10 | 2054.75 | 19.60 | 0.00 | 1.00 | -1017.38 |
| p(Count.Time + Count.Time2 + Noise), N(Elevation + Elevation2 + Latitude + Latitude2) | PZIP | 10 | 2054.80 | 19.64 | 0.00 | 1.00 | -1017.40 |
| p(Date + Noise), N(Elevation + Elevation2 + Latitude + Latitude2) | PZIP | 9 | 2055.43 | 20.27 | 0.00 | 1.00 | -1018.72 |
| p(Count.Time + Count.Time2 + Date + Noise), N(Elevation + Elevation2 + Latitude + Latitude2) | PZIP | 11 | 2055.76 | 20.60 | 0.00 | 1.00 | -1016.88 |

Model selection step 3:

| Candidate model | Distribution | K | AIC | ΔAIC | AIC.Wt | Cum.Wt | LL |
|---|---|---|---|---|---|---|---|
| p(Count.Time + Date + Date2 + Noise + Noise2), N(Elevation + Elevation2 + Latitude + Latitude2) | PZIP | 12 | 2035.16 | 0.00 | 0.64 | 0.64 | -1005.58 |
| p(Count.Time + Date + Date2 + Noise + Noise2 ), N(Latitude + Latitude2) | PZIP | 10 | 2037.14 | 1.98 | 0.24 | 0.88 | -1008.57 |
| p(Count.Time + Date + Date2 + Noise + Noise2 ), N(Elevation + Latitude + Latitude2) | PZIP | 11 | 2038.62 | 3.47 | 0.11 | 0.99 | -1008.31 |
| p(Count.Time + Date + Date2 + Noise + Noise2 ), N(Elevation + Elevation2 + Latitude) | PZIP | 11 | 2043.87 | 8.71 | 0.01 | 1.00 | -1010.93 |
| p(Count.Time + Date + Date2 + Noise + Noise2 ), N(Elevation + Elevation2) | PZIP | 10 | 2051.56 | 16.40 | 0.00 | 1.00 | -1015.78 |
| p(Count.Time + Date + Date2 + Noise + Noise2 ), N(Elevation + Latitude) | PZIP | 10 | 2051.79 | 16.64 | 0.00 | 1.00 | -1015.90 |
| p(Count.Time + Date + Date2 + Noise + Noise2 ), N(Latitude) | PZIP | 9 | 2053.02 | 17.86 | 0.00 | 1.00 | -1017.51 |
| p(Count.Time + Date + Date2 + Noise + Noise2 ), N(Elevation) | PZIP | 9 | 2054.51 | 19.36 | 0.00 | 1.00 | -1018.26 |
