## Supplemental Table 2 for "Accounting for point count ambient noise increases population size estimates"

Supplementary Table S2

| Winter Wren--model selection step 1: | Distribution | K | AIC | ΔAIC | AIC.Wt | Cum.Wt | LL |
|---|---|---|---|---|---|---|---|
| p(Count.Time + Count.Time2 + JDAY + JDAY2 + Noise + Noise2), N(Elevation + Elevation2 + Latitude + Latitude2) | PN | 12 | 3413.06 | 0.00 | 0.58 | 0.58 | -1694.53 |
| p(Count.Time + Count.Time2 + JDAY + JDAY2 + Noise + Noise2), N(Elevation + Elevation2 + Latitude + Latitude2) | NB | 13 | 3415.07 | 2.00 | 0.21 | 0.79 | -1694.53 |
| p(Count.Time + Count.Time2 + JDAY + JDAY2 + Noise + Noise2), N(Elevation + Elevation2 + Latitude + Latitude2) | PZIP | 13 | 3415.07 | 2.01 | 0.21 | 1.00 | -1694.54 |

Model selection step 2:

| Candidate model | Distribution | K | AIC | ΔAIC | AIC.Wt | Cum.Wt | LL |
|---|---|---|---|---|---|---|---|
| p(Count.Time + Count.Time2 + Noise), N(Elevation + Elevation2 + Latitude + Latitude2) | PN | 9 | 3410.05 | 0.00 | 0.26 | 0.26 | -1696.02 |
| p(Count.Time + Count.Time2 + Date + Date2 + Noise), N(Elevation + Elevation2 + Latitude + Latitude2) | PN | 11 | 3411.07 | 1.03 | 0.16 | 0.42 | -1694.54 |
| p(Count.Time + Count.Time2 + Date + Noise), N(Elevation + Elevation2 + Latitude + Latitude2) | PN | 10 | 3411.74 | 1.69 | 0.11 | 0.53 | -1695.87 |
| p(Count.Time + Count.Time2 + Noise + Noise2), N(Elevation + Elevation2 + Latitude + Latitude2) | PN | 10 | 3412.03 | 1.99 | 0.10 | 0.63 | -1696.02 |
| p(Count.Time + Count.Time2 + Date + Date2 + Noise + Noise2), N(Elevation + Elevation2 + Latitude + Latitude2) | PN | 12 | 3413.06 | 3.02 | 0.06 | 0.68 | -1694.53 |
| p(Count.Time + Count.Time2), N(Elevation + Elevation2 + Latitude + Latitude2) | PN | 8 | 3413.12 | 3.07 | 0.06 | 0.74 | -1698.56 |
| p(Count.Time + Count.Time2 + Date + Noise + Noise2), N(Elevation + Elevation2 + Latitude + Latitude2) | PN | 11 | 3413.74 | 3.69 | 0.04 | 0.78 | -1695.87 |
| p(Count.Time + Count.Time2 + Date + Date2), N(Elevation + Elevation2 + Latitude + Latitude2) | PN | 10 | 3413.84 | 3.80 | 0.04 | 0.82 | -1696.92 |
| p(Count.Time + Count.Time2 + Date), N(Elevation + Elevation2 + Latitude + Latitude2) | PN | 9 | 3414.47 | 4.42 | 0.03 | 0.85 | -1698.24 |
| p(Noise), N(Elevation + Elevation2 + Latitude + Latitude2) | PN | 7 | 3414.86 | 4.81 | 0.02 | 0.87 | -1700.43 |
| p(Count.Time + Noise), N(Elevation + Elevation2 + Latitude + Latitude2) | PN | 8 | 3415.15 | 5.10 | 0.02 | 0.89 | -1699.58 |
| p(Count.Time + Date + Date2 + Noise), N(Elevation + Elevation2 + Latitude + Latitude2) | PN | 10 | 3416.20 | 6.16 | 0.01 | 0.90 | -1698.10 |
| p(Date + Date2 + Noise), N(Elevation + Elevation2 + Latitude + Latitude2) | PN | 9 | 3416.26 | 6.22 | 0.01 | 0.92 | -1699.13 |
| p(Date + Noise), N(Elevation + Elevation2 + Latitude + Latitude2) | PN | 8 | 3416.27 | 6.23 | 0.01 | 0.93 | -1700.14 |
| p(Count.Time + Date + Noise), N(Elevation + Elevation2 + Latitude + Latitude2) | PN | 9 | 3416.59 | 6.55 | 0.01 | 0.94 | -1699.30 |
| p(Noise + Noise2), N(Elevation + Elevation2 + Latitude + Latitude2) | PN | 8 | 3416.85 | 6.81 | 0.01 | 0.95 | -1700.43 |
| p(Count.Time), N(Elevation + Elevation2 + Latitude + Latitude2) | PN | 7 | 3417.10 | 7.05 | 0.01 | 0.95 | -1701.55 |
| p(Count.Time + Noise + Noise2), N(Elevation + Elevation2 + Latitude + Latitude2) | PN | 9 | 3417.15 | 7.10 | 0.01 | 0.96 | -1699.58 |
| p(Date + Date2), N(Elevation + Elevation2 + Latitude + Latitude2) | PN | 8 | 3417.54 | 7.50 | 0.01 | 0.97 | -1700.77 |
| p(Date), N(Elevation + Elevation2 + Latitude + Latitude2) | PN | 7 | 3417.55 | 7.50 | 0.01 | 0.97 | -1701.78 |
| p(Count.Time + Date + Date2), N(Elevation + Elevation2 + Latitude + Latitude2) | PN | 9 | 3417.84 | 7.79 | 0.01 | 0.98 | -1699.92 |
| p(Count.Time + Date), N(Elevation + Elevation2 + Latitude + Latitude2) | PN | 8 | 3418.19 | 8.15 | 0.00 | 0.98 | -1701.10 |
| p(Count.Time + Date + Date2 + Noise + Noise2), N(Elevation + Elevation2 + Latitude + Latitude2) | PN | 11 | 3418.20 | 8.15 | 0.00 | 0.99 | -1698.10 |
| p(Date + Noise + Noise2), N(Elevation + Elevation2 + Latitude + Latitude2) | PN | 9 | 3418.24 | 8.20 | 0.00 | 0.99 | -1700.12 |
| p(Date + Date2 + Noise + Noise2), N(Elevation + Elevation2 + Latitude + Latitude2) | PN | 10 | 3418.25 | 8.20 | 0.00 | 1.00 | -1699.12 |
| p(Count.Time + Date + Noise + Noise2), N(Elevation + Elevation2 + Latitude + Latitude2) | PN | 10 | 3418.58 | 8.54 | 0.00 | 1.00 | -1699.29 |

Model selection step 3:

| Candidate model | Distribution | K | AIC | ΔAIC | AIC.Wt | Cum.Wt | LL |
|---|---|---|---|---|---|---|---|
| p(Count.Time + Count.Time2 + Noise), N(Elevation + Elevation2) | PN | 7 | 3406.63 | 0.00 | 0.56 | 0.56 | -1696.31 |
| p(Count.Time + Count.Time2 + Noise), N(Elevation + Elevation2 + Latitude) | PN | 8 | 3408.26 | 1.63 | 0.25 | 0.81 | -1696.13 |
| p(Count.Time + Count.Time2 + Noise), N(Elevation + Elevation2 + Latitude + Latitude2) | PN | 9 | 3410.05 | 3.42 | 0.10 | 0.91 | -1696.02 |
| p(Count.Time + Count.Time2 + Noise), N(Latitude) | PN | 6 | 3412.05 | 5.42 | 0.04 | 0.95 | -1700.02 |
| p(Count.Time + Count.Time2 + Noise), N(Elevation) | PN | 6 | 3413.86 | 7.23 | 0.02 | 0.97 | -1700.93 |
| p(Count.Time + Count.Time2 + Noise), N(Elevation + Latitude) | PN | 7 | 3413.92 | 7.29 | 0.01 | 0.98 | -1699.96 |
| p(Count.Time + Count.Time2 + Noise), N(Latitude + Latitude2) | PN | 7 | 3414.04 | 7.41 | 0.01 | 0.99 | -1700.02 |
| p(Count.Time + Count.Time2 + Noise), N(Elevation + Latitude + Latitude2) | PN | 8 | 3415.90 | 9.27 | 0.01 | 1.00 | -1699.95 |
