## Supplemental Table 3 for "Accounting for point count ambient noise increases population size estimates"

Supplementary Table S3

| Candidate model | Distribution | K | AIC | ΔAIC | AIC.Wt | Cum.Wt | LL |
|---|---|---|---|---|---|---|---|
| Blackpoll Warbler--model selection step 1: | | | | | | | |
| p(Count.Time + Count.Time2 + Date + Date2 + Noise + Noise2), N(Elevation + Elevation2 + Latitude + Latitude2) | PN | 12 | 3094.780 | 0.000 | 0.580 | 0.580 | -1535.390 |
| p(Count.Time + Count.Time2 + Date + Date2 + Noise + Noise2), N(Elevation + Elevation2 + Latitude + Latitude2) | NB | 13 | 3096.780 | 2.000 | 0.210 | 0.790 | -1535.390 |
| p(Count.Time + Count.Time2 + Date + Date2 + Noise + Noise2), N(Elevation + Elevation2 + Latitude + Latitude2) | PZIP | 13 | 3096.800 | 2.020 | 0.210 | 1.000 | -1535.400 |

| Candidate model | Distribution | K | AIC | ΔAIC | AIC.Wt | Cum.Wt | LL |
|---|---|---|---|---|---|---|---|
| Model selection step 2: | | | | | | | |
| p(Date + Noise + Noise2), N(Elevation + Elevation2 + Latitude + Latitude2) | PN | 9 | 3088.90 | 0.00 | 0.37 | 0.37 | -1535.45 |
| p(Noise + Noise2), N(Elevation + Elevation2 + Latitude + Latitude2) | PN | 8 | 3090.75 | 1.85 | 0.15 | 0.51 | -1537.38 |
| p(Count.Time + Date + Noise + Noise2), N(Elevation + Elevation2 + Latitude + Latitude2) | PN | 10 | 3090.82 | 1.92 | 0.14 | 0.65 | -1535.41 |
| p(Date + Date2 + Noise + Noise2), N(Elevation + Elevation2 + Latitude + Latitude2) | PN | 10 | 3090.87 | 1.97 | 0.14 | 0.79 | -1535.44 |
| p(Count.Time + Noise + Noise2), N(Elevation + Elevation2 + Latitude + Latitude2) | PN | 9 | 3092.61 | 3.70 | 0.06 | 0.85 | -1537.30 |
| p(Count.Time + Date + Date2 + Noise + Noise2), N(Elevation + Elevation2 + Latitude + Latitude2) | PN | 11 | 3092.80 | 3.89 | 0.05 | 0.90 | -1535.40 |
| p(Count.Time + Count.Time2 + Date + Noise + Noise2), N(Elevation + Elevation2 + Latitude + Latitude2) | PN | 11 | 3092.81 | 3.91 | 0.05 | 0.95 | -1535.40 |
| p(Count.Time + Count.Time2 + Noise + Noise2), N(Elevation + Elevation2 + Latitude + Latitude2) | PN | 10 | 3094.58 | 5.68 | 0.02 | 0.97 | -1537.29 |
| p(Count.Time + Count.Time2 + Date + Date2 + Noise + Noise2), N(Elevation + Elevation2 + Latitude + Latitude2) | PN | 12 | 3094.78 | 5.88 | 0.02 | 0.99 | -1535.39 |
| p(Date), N(Elevation + Elevation2 + Latitude + Latitude2) | PN | 7 | 3099.03 | 10.13 | 0.00 | 0.99 | -1542.51 |
| p(Count.Time), N(Elevation + Elevation2 + Latitude + Latitude2) | PN | 7 | 3100.80 | 11.90 | 0.00 | 0.99 | -1543.40 |
| p(Count.Time + Date), N(Elevation + Elevation2 + Latitude + Latitude2) | PN | 8 | 3100.84 | 11.94 | 0.00 | 0.99 | -1542.42 |
| p(Date + Noise), N(Elevation + Elevation2 + Latitude + Latitude2) | PN | 8 | 3100.98 | 12.08 | 0.00 | 1.00 | -1542.49 |
| p(Noise), N(Elevation + Elevation2 + Latitude + Latitude2) | PN | 7 | 3101.01 | 12.10 | 0.00 | 1.00 | -1543.50 |
| p(Date + Date2), N(Elevation + Elevation2 + Latitude + Latitude2) | PN | 8 | 3101.03 | 12.12 | 0.00 | 1.00 | -1542.51 |
| p(Count.Time + Count.Time2), N(Elevation + Elevation2 + Latitude + Latitude2) | PN | 8 | 3102.73 | 13.83 | 0.00 | 1.00 | -1543.37 |
| p(Count.Time + Noise), N(Elevation + Elevation2 + Latitude + Latitude2) | PN | 8 | 3102.76 | 13.85 | 0.00 | 1.00 | -1543.38 |
| p(Count.Time + Count.Time2 + Date), N(Elevation + Elevation2 + Latitude) | PN | 9 | 3102.77 | 13.87 | 0.00 | 1.00 | -1542.39 |
| p(Count.Time + Date + Noise), N(Elevation + Elevation2 + Latitude) | PN | 9 | 3102.78 | 13.88 | 0.00 | 1.00 | -1542.39 |
| p(Count.Time + Date + Date2), N(Elevation + Elevation2 + Latitude) | PN | 9 | 3102.84 | 13.94 | 0.00 | 1.00 | -1542.42 |
| p(Date + Date2 + Noise), N(Elevation + Elevation2 + Latitude) | PN | 9 | 3102.97 | 14.07 | 0.00 | 1.00 | -1542.49 |
| p(Count.Time + Count.Time2 + Noise), N(Elevation + Elevation2 + Latitude) | PN | 9 | 3104.68 | 15.78 | 0.00 | 1.00 | -1543.34 |
| p(Count.Time + Count.Time2 + Date + Noise), N(Elevation + Elevation2 + Latitude) | PN | 10 | 3104.71 | 15.81 | 0.00 | 1.00 | -1542.36 |
| p(Count.Time + Count.Time2 + Date + Date2), N(Elevation + Elevation2 + Latitude) | PN | 10 | 3104.77 | 15.87 | 0.00 | 1.00 | -1542.39 |
| p(Count.Time + Date + Date2 + Noise), N(Elevation + Elevation2 + Latitude) | PN | 10 | 3104.78 | 15.88 | 0.00 | 1.00 | -1542.39 |
| p(Count.Time + Count.Time2 + Date + Date2 + Noise), N(Elevation + Elevation2 + Latitude) | PN | 11 | 3106.71 | 17.81 | 0.00 | 1.00 | -1542.36 |

| Candidate model | Distribution | K | AIC | ΔAIC | AIC.Wt | Cum.Wt | LL |
|---|---|---|---|---|---|---|---|
| Model selection step 3: | | | | | | | |
| Date + Noise + Noise2), N(Elevation + Elevation2 + Latitude) | PN | 8 | 3088.44 | 0.00 | 0.55 | 0.55 | -1536.22 |
| Date + Noise + Noise2), N(Elevation + Elevation2 + Latitude + Latitude2) | PN | 9 | 3088.90 | 0.46 | 0.44 | 0.99 | -1535.45 |
| Date + Noise + Noise2), N(Elevation + Elevation2) | PN | 7 | 3096.44 | 8.00 | 0.01 | 1.00 | -1541.22 |
| Date + Noise + Noise2), N(Elevation + Latitude) | PN | 7 | 3105.92 | 17.48 | 0.00 | 1.00 | -1545.96 |
| Date + Noise + Noise2), N(Elevation + Latitude + Latitude2) | PN | 8 | 3107.60 | 19.16 | 0.00 | 1.00 | -1545.80 |
| Date + Noise + Noise2), N(Elevation) | PN | 6 | 3107.76 | 19.32 | 0.00 | 1.00 | -1547.88 |
| Date + Noise + Noise2), N(Latitude) | PN | 6 | 3151.25 | 62.81 | 0.00 | 1.00 | -1569.63 |
| Date + Noise + Noise2), N(Latitude + Latitude2) | PN | 7 | 3151.94 | 63.49 | 0.00 | 1.00 | -1568.97 |
