## Supplemental Table 4 for "Accounting for point count ambient noise increases population size estimates"

Supplementary Table S4

White-throated Sparrow--model selection step 1:

| Candidate model | Distribution | K | AIC | ΔAIC | AIC.Wt | Cum.Wt | LL |
|---|---|---|---|---|---|---|---|
| p(Count.Time + Count.Time2 + JDAY + JDAY2 + Noise + Noise2), N(Elevation + Elevation2 + Latitude + Latitude2) | PZIP | 13 | 2982.54 | 0.00 | 0.63 | 0.63 | -1478.27 |
| p(Count.Time + Count.Time2 + JDAY + JDAY2 + Noise + Noise2), N(Elevation + Elevation2 + Latitude + Latitude2) | NB | 13 | 2983.57 | 1.03 | 0.37 | 1.00 | -1478.79 |
| p(Count.Time + Count.Time2 + JDAY + JDAY2 + Noise + Noise2), N(Elevation + Elevation2 + Latitude + Latitude2) | PN | 12 | 3020.07 | 37.52 | 0.00 | 1.00 | -1498.03 |

Model selection step 2:

| Candidate model | Distribution | K | AIC | ΔAIC | AIC.Wt | Cum.Wt | LL |
|---|---|---|---|---|---|---|---|
| p(Count.Time + Count.Time2 + JDAY + JDAY2 + Noise + Noise2), N(Elevation + Elevation2 + Latitude + Latitude2) | PZIP | 13 | 2982.54 | 0.00 | 0.62 | 0.62 | -1478.27 |
| p(Count.Time + Count.Time2 + Date + Date2 + Noise), N(Elevation + Elevation2 + Latitude + Latitude2) | PZIP | 12 | 2985.06 | 2.52 | 0.18 | 0.80 | -1480.53 |
| p(Count.Time + Count.Time2 + Date + Date2), N(Elevation + Elevation2 + Latitude + Latitude2) | PZIP | 11 | 2986.31 | 3.77 | 0.09 | 0.89 | -1482.16 |
| p(Count.Time + Count.Time2 + Date + Noise + Noise2), N(Elevation + Elevation2 + Latitude + Latitude2) | PZIP | 12 | 2986.71 | 4.17 | 0.08 | 0.97 | -1481.36 |
| p(Count.Time + Count.Time2 + Date + Noise), N(Elevation + Elevation2 + Latitude + Latitude2) | PZIP | 11 | 2989.82 | 7.28 | 0.02 | 0.98 | -1483.91 |
| p(Count.Time + Count.Time2 + Date), N(Elevation + Elevation2 + Latitude + Latitude2) | PZIP | 10 | 2991.74 | 9.19 | 0.01 | 0.99 | -1485.87 |
| p(Count.Time + Count.Time2 + Noise + Noise2), N(Elevation + Elevation2 + Latitude + Latitude2) | PZIP | 11 | 2992.78 | 10.23 | 0.00 | 0.99 | -1485.39 |
| p(Count.Time + Date + Date2 + Noise + Noise2), N(Elevation + Elevation2 + Latitude + Latitude2) | PZIP | 12 | 2993.00 | 10.46 | 0.00 | 1.00 | -1484.50 |
| p(Count.Time + Date + Noise + Noise2), N(Elevation + Elevation2 + Latitude + Latitude2) | PZIP | 11 | 2995.48 | 12.94 | 0.00 | 1.00 | -1486.74 |
| p(Count.Time + Count.Time2 + Noise), N(Elevation + Elevation2 + Latitude + Latitude2) | PZIP | 10 | 2997.22 | 14.67 | 0.00 | 1.00 | -1488.61 |
| p(Count.Time + Count.Time2), N(Elevation + Elevation2 + Latitude + Latitude2) | PZIP | 9 | 2997.49 | 14.94 | 0.00 | 1.00 | -1489.74 |
| p(Count.Time + Date + Date2 + Noise), N(Elevation + Elevation2 + Latitude + Latitude2) | PZIP | 11 | 2998.96 | 16.42 | 0.00 | 1.00 | -1488.48 |
| p(Count.Time + Date + Date2), N(Elevation + Elevation2 + Latitude + Latitude2) | PZIP | 10 | 3000.36 | 17.81 | 0.00 | 1.00 | -1490.18 |
| p(Date + Date2 + Noise + Noise2), N(Elevation + Elevation2 + Latitude + Latitude2) | PZIP | 11 | 3000.90 | 18.35 | 0.00 | 1.00 | -1489.45 |
| p(Count.Time + Noise + Noise2), N(Elevation + Elevation2 + Latitude + Latitude2) | PZIP | 10 | 3001.72 | 19.18 | 0.00 | 1.00 | -1490.86 |
| p(Count.Time + Date + Noise), N(Elevation + Elevation2 + Latitude + Latitude2) | PZIP | 10 | 3001.81 | 19.26 | 0.00 | 1.00 | -1490.90 |
| p(Date + Noise + Noise2), N(Elevation + Elevation2 + Latitude + Latitude2) | PZIP | 10 | 3001.86 | 19.32 | 0.00 | 1.00 | -1490.93 |
| p(Count.Time + Date), N(Elevation + Elevation2 + Latitude + Latitude2) | PZIP | 9 | 3003.74 | 21.20 | 0.00 | 1.00 | -1492.87 |
| p(Noise + Noise2), N(Elevation + Elevation2 + Latitude + Latitude2) | PZIP | 9 | 3007.03 | 24.49 | 0.00 | 1.00 | -1494.52 |
| p(Date + Date2 + Noise), N(Elevation + Elevation2 + Latitude + Latitude2) | PZIP | 10 | 3008.99 | 26.45 | 0.00 | 1.00 | -1494.49 |
| p(Count.Time + Noise), N(Elevation + Elevation2 + Latitude + Latitude2) | PZIP | 9 | 3009.85 | 27.30 | 0.00 | 1.00 | -1495.92 |
| p(Date + Date2), N(Elevation + Elevation2 + Latitude + Latitude2) | PZIP | 9 | 3010.07 | 27.53 | 0.00 | 1.00 | -1496.03 |
| p(Count.Time), N(Elevation + Elevation2 + Latitude + Latitude2) | PZIP | 8 | 3010.12 | 27.57 | 0.00 | 1.00 | -1497.06 |
| p(Date + Noise), N(Elevation + Elevation2 + Latitude + Latitude2) | PZIP | 9 | 3010.12 | 27.58 | 0.00 | 1.00 | -1496.06 |
| p(Date), N(Elevation + Elevation2 + Latitude + Latitude2) | PZIP | 8 | 3011.75 | 29.20 | 0.00 | 1.00 | -1497.87 |
| p(Noise), N(Elevation + Elevation2 + Latitude + Latitude2) | PZIP | 8 | 3016.67 | 34.12 | 0.00 | 1.00 | -1500.33 |

Model selection step 3:

| Candidate model | Distribution | K | AIC | ΔAIC | AIC.Wt | Cum.Wt | LL |
|---|---|---|---|---|---|---|---|
| p(Count.Time + Count.Time2 + Date + Date2 + Noise + Noise2), N(Elevation + Latitude) | PZIP | 11 | 2979.63 | 0.00 | 0.44 | 0.44 | -1478.81 |
| p(Count.Time + Count.Time2 + Date + Date2 + Noise + Noise2), N(Elevation + Elevation2 + Latitude) | PZIP | 12 | 2980.69 | 1.06 | 0.26 | 0.70 | -1478.34 |
| p(Count.Time + Count.Time2 + Date + Date2 + Noise + Noise2), N(Elevation + Latitude + Latitude2) | PZIP | 12 | 2981.34 | 1.72 | 0.19 | 0.89 | -1478.67 |
| p(Count.Time + Count.Time2 + Date + Date2 + Noise + Noise2), N(Elevation + Elevation2 + Latitude + Latitude2) | PZIP | 13 | 2982.54 | 2.92 | 0.10 | 0.99 | -1478.27 |
| p(Count.Time + Count.Time2 + Date + Date2 + Noise + Noise2), N(Latitude) | PZIP | 10 | 2987.88 | 8.25 | 0.01 | 1.00 | -1483.94 |
| p(Count.Time + Count.Time2 + Date + Date2 + Noise + Noise2), N(Latitude + Latitude2) | PZIP | 11 | 2989.77 | 10.15 | 0.00 | 1.00 | -1483.89 |
| p(Count.Time + Count.Time2 + Date + Date2 + Noise + Noise2), N(Elevation + Elevation2) | PZIP | 11 | 3016.33 | 36.70 | 0.00 | 1.00 | -1497.17 |
| p(Count.Time + Count.Time2 + Date + Date2 + Noise + Noise2), N(Elevation) | PZIP | 10 | 3022.29 | 42.67 | 0.00 | 1.00 | -1501.15 |
