## Supplemental Table 5 for "Accounting for point count ambient noise increases population size estimates"

Supplementary Table S5

| Yellow-bellied Flycatcher: Parameters | β | SD | LCRI | UCRI |
|---|---|---|---|---|
| p: Intercept | 0.593686 | 0.039806 | 0.513541 | 0.670054 |
| p: Count.Time | -0.21433 | 0.100423 | -0.4173 | -0.01668 |
| p: Count.Time2 | N/A | | | |
| p: Date | -0.12048 | 0.101302 | -0.31861 | 0.078124 |
| p: Date2 | -0.36168 | 0.095274 | -0.5547 | -0.18168 |
| p: Noise | 0.449482 | 0.13103 | 0.197988 | 0.714705 |
| p: Noise2 | -0.21435 | 0.058995 | -0.32691 | -0.09937 |
| N: Intercept | -0.03113 | 0.151352 | -0.34094 | 0.254505 |
| N: Elevation | 0.162869 | 0.097786 | -0.02477 | 0.358233 |
| N: Elevation2 | -0.15577 | 0.068595 | -0.29161 | -0.02811 |
| N: Latitude | 0.188763 | 0.117449 | -0.04166 | 0.423823 |
| N: Latitude2 | -0.19912 | 0.065539 | -0.3353 | -0.07693 |
| Φ: Proportion of suitable sites | 0.699737 | 0.075243 | 0.569597 | 0.860607 |
| Derived: totalN | 295.2427 | 14.61747 | 270 | 327 |
| Derived: mean.detection | 0.459042 | 0.024233 | 0.410592 | 0.504415 |
