## Supplemental Table 6 for "Accounting for point count ambient noise increases population size estimates"

Supplementary Table S6

| Winter Wren: Parameters | β | SD | LCRI | UCRI |
|---|---|---|---|---|
| p: Intercept | 0.60299 | 0.081607 | 0.436027 | 0.759602 |
| p: Count.Time | -0.08682 | 0.066125 | -0.21632 | 0.038733 |
| p: Count.Time2 | -0.15805 | 0.060604 | -0.2746 | -0.03492 |
| p: Date | N/A | | | |
| p: Date2 | N/A | | | |
| p: Noise | -0.13015 | 0.058256 | -0.24826 | -0.01868 |
| p: Noise2 | N/A | | | |
| N: Intercept | 0.076607 | 0.057885 | -0.03693 | 0.187684 |
| N: Elevation | 0.060139 | 0.038026 | 0.003738 | 0.145915 |
| N: Elevation2 | -0.1226 | 0.039912 | -0.20089 | -0.04602 |
| N: Latitude | N/A | | | |
| N: Latitude2 | N/A | | | |
| Φ: Proportion of suitable sites | N/A | | | |
| Derived: totalN | 539.581 | 9.941045 | 522 | 561 |
| Derived: mean.detection | 0.608236 | 0.015136 | 0.577453 | 0.637265 |
