## Supplemental Table 7 for "Accounting for point count ambient noise increases population size estimates"

Supplementary Table S7

| Blackpoll Warbler: Parameters | β | SD | LCRI | UCRI |
|---|---|---|---|---|
| p: Intercept | 0.361508 | 0.080191 | 0.204589 | 0.513561 |
| p: Count.Time | N/A | | | |
| p: Count.Time2 | N/A | | | |
| p: Date | -0.1439 | 0.070654 | -0.27964 | -0.006 |
| p: Date2 | N/A | | | |
| p: Noise | 0.295426 | 0.097065 | 0.104405 | 0.487056 |
| p: Noise2 | -0.16713 | 0.044114 | -0.25111 | -0.08183 |
| N: Intercept | -0.04537 | 0.06233 | -0.16785 | 0.076959 |
| N: Elevation | 0.534772 | 0.069143 | 0.396995 | 0.670675 |
| N: Elevation2 | -0.19881 | 0.046821 | -0.29529 | -0.11338 |
| N: Latitude | 0.179003 | 0.056735 | 0.070843 | 0.290696 |
| N: Latitude2 | N/A | | | |
| Φ: Proportion of suitable sites | N/A | | | |
| Derived: totalN | 481.9753 | 11.98766 | 461 | 508 |
| Derived: mean.detection | 0.548014 | 0.017047 | 0.514616 | 0.580037 |
