## Supplemental Table 8 for "Accounting for point count ambient noise increases population size estimates"

Supplementary Table S8

| White-throated Sparrow: Parameters | β | SD | LCRI | UCRI |
|---|---|---|---|---|
| p: Intercept | -0.35124 | 0.17655 | -0.69724 | -0.008 |
| p: Count.Time | -0.21005 | 0.069403 | -0.35061 | -0.07742 |
| p: Count.Time2 | 0.196302 | 0.056638 | 0.084426 | 0.306803 |
| p: Date | 0.146839 | 0.067623 | 0.015372 | 0.280113 |
| p: Date2 | 0.134114 | 0.056741 | 0.025027 | 0.249203 |
| p: Noise | 0.019454 | 0.108378 | -0.20281 | 0.230953 |
| p: Noise2 | -0.11744 | 0.055032 | -0.22461 | -0.00863 |
| N: Intercept | 0.204408 | 0.086157 | 0.037438 | 0.370503 |
| N: Elevation | 0.195181 | 0.061637 | 0.071638 | 0.315627 |
| N: Elevation2 | N/A | | | |
| N: Latitude | 0.509897 | 0.080975 | 0.357897 | 0.6689 |
| N: Latitude2 | N/A | | | |
| Φ: Proportion of suitable sites | 0.706859 | 0.039795 | 0.63166 | 0.787117 |
| Derived: totalN | 525.5297 | 21.33226 | 489 | 572 |
| Derived: mean.detection | 0.467947 | 0.022466 | 0.4238 | 0.510407 |
