## Supplemental Table 9 for "Accounting for point count ambient noise increases population size estimates"

Supplementary Table S9

| Parameter | Univariate | | Quadratic | |
|---|---|---|---|---|
| | Mean | SD | Mean | SD |
| Date | -0.1439 | 0.070654 | 0.013182 | 0.159008 |
| Date2 | N/A | | -0.11378 | 0.260019 |
| Count.Time | -0.21433 | 0.100423 | -0.14843 | 0.091605 |
| Count.Time2 | N/A | | 0.019125 | 0.186646 |
| Noise | -0.13015 | 0.058256 | 0.254788 | 0.210774 |
| Noise2 | N/A | | -0.16631 | 0.066208 |
| Elevation | 0.195181 | 0.061637 | 0.252593 | 0.21642 |
| Elevation2 | N/A | | -0.15906 | 0.06167 |
| Latitude | 0.34445 | 0.179623 | 0.188763 | 0.117449 |
| Latitude2 | N/A | | -0.19912 | 0.065539 |
